# DNA breaks-mediated cost reveals RNase HI as a new target for selectively eliminating antibiotic resistance

**DOI:** 10.1101/756767

**Authors:** Roberto Balbontín, Nelson Frazão, Isabel Gordo

**Affiliations:** Instituto Gulbenkian de Ciência; Oeiras, 2780-156; Portugal

**Keywords:** antibiotic resistance, fitness cost, DNA breaks, RNase HI targeting, repurposed drug

## Abstract

Antibiotic resistance often generates a fitness cost to bacteria in drug-free environments. Understanding the causes of the cost is considered the Holy Grail in the antibiotic resistance field, as it is the main determinant of the prevalence of resistances upon reducing antibiotics use. We show that DNA breaks can explain most of the variation in the cost of resistances common in pathogens. Here we demonstrate that targeting the RNase that degrades R-loops, which cause DNA breaks, exacerbates the cost of resistance. Consequently, lack of RNase HI function drives resistant clones to extinction in populations with high initial frequency of resistance, both in laboratory conditions and in a mouse model of gut colonization. Thus, RNase HI provides a target specific against resistant bacteria, which we validate using a repurposed drug. In summary, we revealed key mechanisms underlying the cost of antibiotic resistance that can be exploited to specifically eliminate resistant bacteria.

## Introduction

Antibiotic resistance entails a large health and economic burden worldwide (1). Its maintenance and spread in bacterial populations depend on the rate at which resistance is acquired and on its effect on bacterial fitness. This effect is typically deleterious, resulting in the so-called cost of resistance. The cost of resistance is influenced by the environment, by interactions between the resistances and the genetic background in which they appear (epistasis), and by subsequent acquisition of mutations compensating for fitness defects (compensatory evolution) (2). Importantly, the magnitude of the cost is the main biological parameter influencing the fate of resistances upon reducing antibiotic use (3). Despite its importance, the causes of the cost are far from being completely understood (4), and the identification of these causes has become the “Holy Grail” in the antibiotic resistance field.

Resistance mutations often map to genes encoding proteins targeted by antibiotics. These target proteins are typically involved in essential functions, such as transcription, translation, DNA replication or cell wall biosynthesis. Resistance mutations cause alterations in the structure of the target protein, rendering it insensitive to the drug, but often adversely affecting its function (2,3,5). These pleiotropic effects hold the key to manipulating resistance levels in bacterial populations (3). Rifampicin and streptomycin resistance mutations (Rif^R^ and Str^R^), common in pathogenic bacteria (6–8), map to the genes *rpoB* and *rpsL*, encoding the *β’* subunit of the RNA polymerase and the protein S12 of the 30S ribosomal subunit, respectively. Rif^R^ and Str^R^ mutations are representative examples of resistances to antibiotics targeting transcription and translation, respectively. Rif^R^ mutations show different costs (9, 10) and cause alterations in the rates of transcription initiation, elongation, slippage or termination (9,11–23). Likewise, most Str^R^ mutations also cause a cost (24–26) and affect translation fidelity and processivity (24,26–34). Thus, at present, the costs of Rif^R^ and Str^R^ mutations are thought to be caused by defects in protein synthesis, either at a global cellular level (35–37) or limited to specific functions or regulons (22,24,38,39).

We recently showed that the fitness cost of double resistance (Rif^R^ Str^R^) can be reduced via overexpression of *nusG* or *nusE* (40), which encode the proteins that physically connect the RNA polymerase and the ribosome (41). This suggests that the reduction of the cost can be achieved by reinforcing the coupling between transcription and translation, besides the previously described partial recovery in protein biosynthesis (3). Coupling between transcription and translation is known to prevent spontaneous backtracking of the RNA polymerase which, if exacerbated, can cause a series of molecular events that ultimately lead to the generation of double-strand DNA breaks (42–46). This led us to hypothesize that cells carrying different Rif^R^ and Str^R^ mutations have higher number of DNA breaks, which greatly contribute to their cost.

In this study, we show that DNA breaks caused by resistance mutations are major contributors to their fitness cost, explaining 73% of its variation across resistant genotypes. The involvement of R-loops in the generation of DNA breaks allowed us to identify RNase HI as an important determinant of the cost of resistance, which we validate using a repurposed drug. We further show that targeting RNase HI is a plausible strategy for the selective elimination of resistant bacteria, as lack of RNase HI leads to extinction of resistant clones when competing against sensitive bacteria in laboratory conditions, abolishing compensatory evolution. Importantly, eradication of resistant bacteria lacking RNase HI is remarkably efficient in the mammalian gut as well. Altogether, we reveal an unexpected cause of the cost of antibiotic resistance, which can be exploited to selectively eliminate resistant bacteria in polymorphic populations.

## Results

### DNA breaks explain variation in the fitness cost of resistance

In order to test if Rif^R^ and Str^R^ mutations generate DNA breaks, and whether DNA breaks contribute to the cost of resistance, we simultaneously measured competitive fitness and activation of the SOS response, a well-known proxy specific for the occurrence of DNA breaks (47, 48). We carried out these assays in sensitive *Escherichia coli*, Str^R^ strains (RpsL^K43N^, RpsL^K43T^, and RpsL^K43R^), Rif^R^ strains (RpoB^H526L^, RpoB^H526Y^, and RpoB^S531F^), and double resistant mutants carrying the nine possible combinations of these resistance alleles. Remarkably, fourteen out of the fifteen resistant strains show increased SOS activation (Figure 1A), demonstrating that resistance mutations indeed cause DNA breaks. Moreover, the SOS induction strongly correlates with the cost of resistance, explaining 73% of its variation (Figure 1B), suggesting that DNA breaks are key contributors to the cost of resistance. To independently confirm the occurrence of DNA breaks in resistant bacteria, we used a system which permits direct visualization of double-stranded DNA ends by fluorescence microscopy (49). We combined this system with the SOS reporter, and analyzed a subset of resistant mutants and sensitive bacteria. As expected, this corroborated that resistance mutations indeed cause increased number of DNA breaks (Table 1, Figure S1). We then asked if resistance mutations involving a different mechanism which could lead to perturbations of transcription-translation coupling also generate DNA breaks. The commonly used antibiotic erythromycin targets the 50S ribosomal subunit, affecting translation and its coupling with transcription (50). Erythromycin resistance mutations (Erm^R^) mapping to the genes *rplD* and *rplV* (encoding the 50S ribosomal subunit proteins L4 and L22, respectively) are known to reduce translation elongation rate (51, 52), likely affecting transcription-translation coupling. We isolated Erm^R^ clones carrying either RplD^G66R^ or RplV^Δ(82-84)^ mutations and found that both mutants show fitness cost and increased SOS (Figure 1C), demonstrating that other resistance mutations, with a different mechanistic basis but affecting the same process (transcription-translation coupling) also cause DNA breaks.

**Figure 1.**
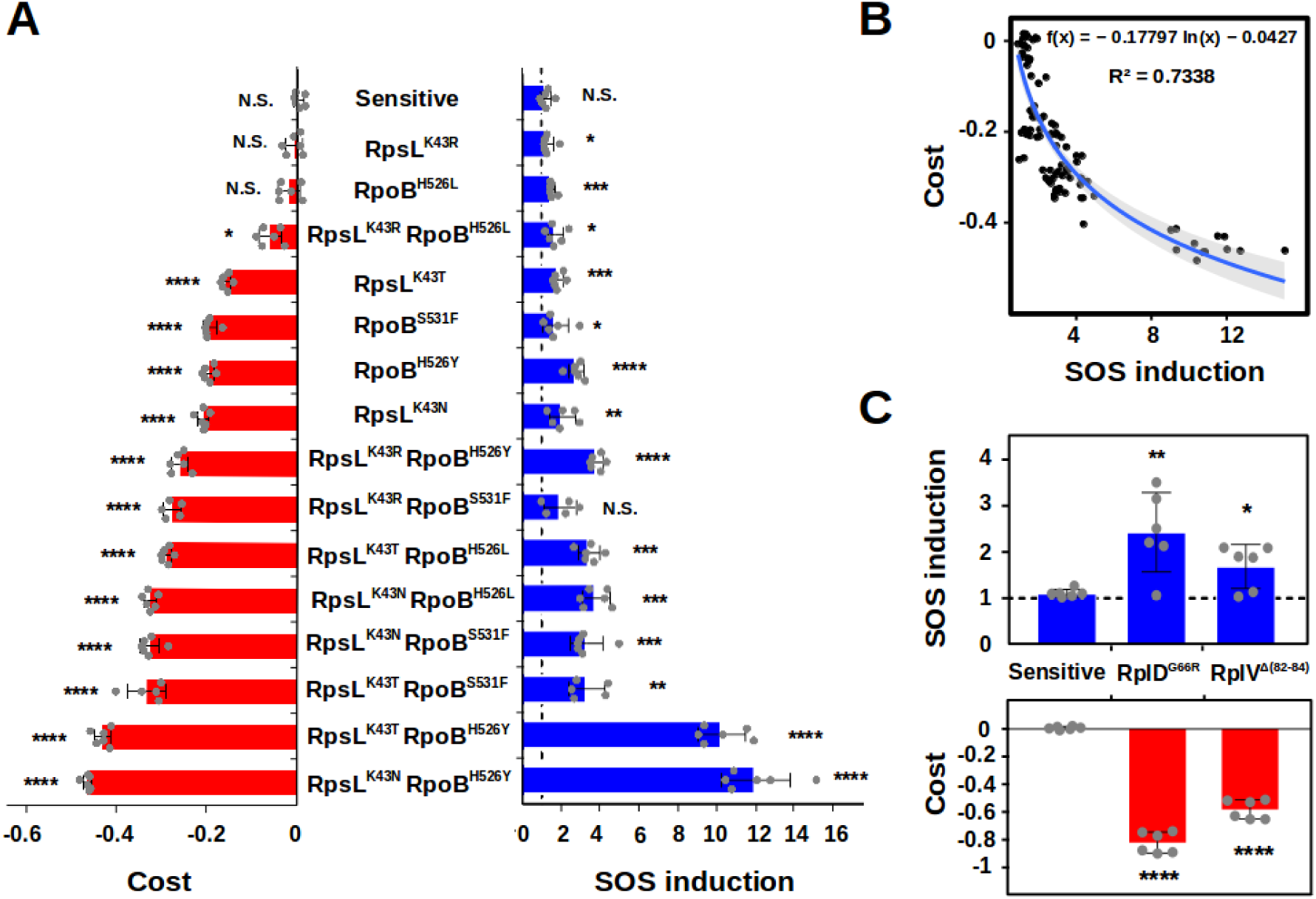
DNA breaks explain variation in the fitness cost of resistance. **A.** Fitness cost (red bars) and SOS induction (blue bars) of sensitive bacteria and Str^R^ and Rif^R^ mutants in LB broth at 4 hours. The strains are ordered from lower to higher fitness cost (top to bottom). The dashed line indicates no SOS induction. Error bars represent mean ± standard deviation of independent biological replicates (n≥5). N.S. non-significant; \**P* < 0.05; \*\**P* < 0.01; \*\*\**P* < 0.001; \*\*\*\**P* < 0.0001 (two-tailed Student’s *t* test). **B.** Correlation between the fitness cost (y axis) and the SOS induction (x axis) representing all the data points from A. The blue line represents the logarithmic regression line, and the grey area represents the 95% confidence interval. **C.** SOS induction (blue bars) and fitness cost (red bars) of sensitive bacteria and Erm^R^ mutants in LB broth at 4 hours. The dashed line indicates no SOS induction. Error bars represent mean ± standard deviation of independent biological replicates (n=6). N.S. non-significant; \**P* < 0.05; \*\**P* < 0.01; \*\*\**P* < 0.001; \*\*\*\**P* < 0.0001 (two-tailed Student’s *t* test).

**Table 1.**
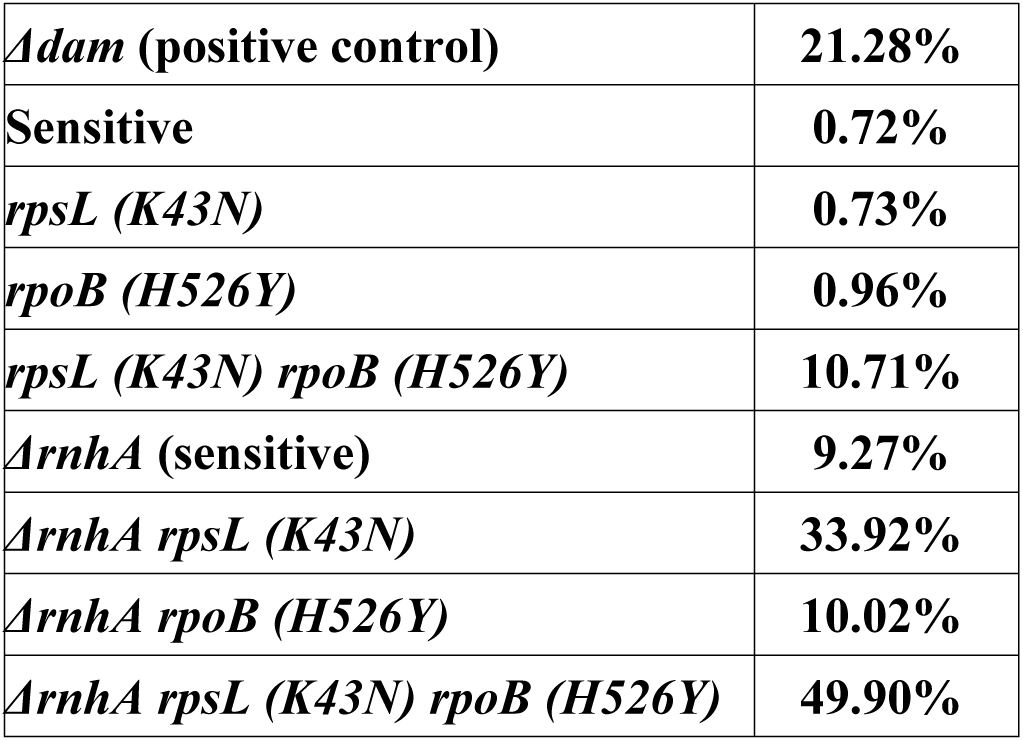
Percentage of cells showing DNA breaks (Gam-GFP foci) in sensitive and resistant bacteria either in wild-type or ΔrnhA backgrounds. Except in the *Δdam* positive control (in which over 100 cells sufficed to provide an illustrative example), at least 1000 cells per group were analyzed. See also Figures S1 and S2.

### Compensatory evolution cause reduction of DNA breaks

We then hypothesized that, if DNA breaks are key contributors to the fitness cost of resistance, compensatory evolution of resistant strains should result in a reduction of DNA breaks. To test this, we compared the cost and SOS induction in the RpsL^K43T^ RpoB^H526Y^ double mutant and in an isogenic strain additionally carrying the most prevalent compensatory mutation found in our previous study: RpoC^Q1126K^ (40). As hypothesized, both cost and SOS induction are greatly reduced in the compensated strain (Figure 2A), confirming that DNA breaks are targeted by compensatory evolution. In order to test if this is general, we analyzed nine compensated clones from three independent populations of the RpsL^K43T^ RpoB^H526Y^ double mutant propagated for fifteen days in the absence of antibiotics. As expected, the costs are smaller in the evolved strains and, as hypothesized, all the nine compensated clones show decreased SOS induction compared to their resistant ancestor (Figure 2B). This demonstrates that compensatory evolution widely targets mechanisms that reduce DNA breaks and further reinforces the notion that DNA breaks are major contributors to the cost of antibiotic resistance.

**Figure 2.**
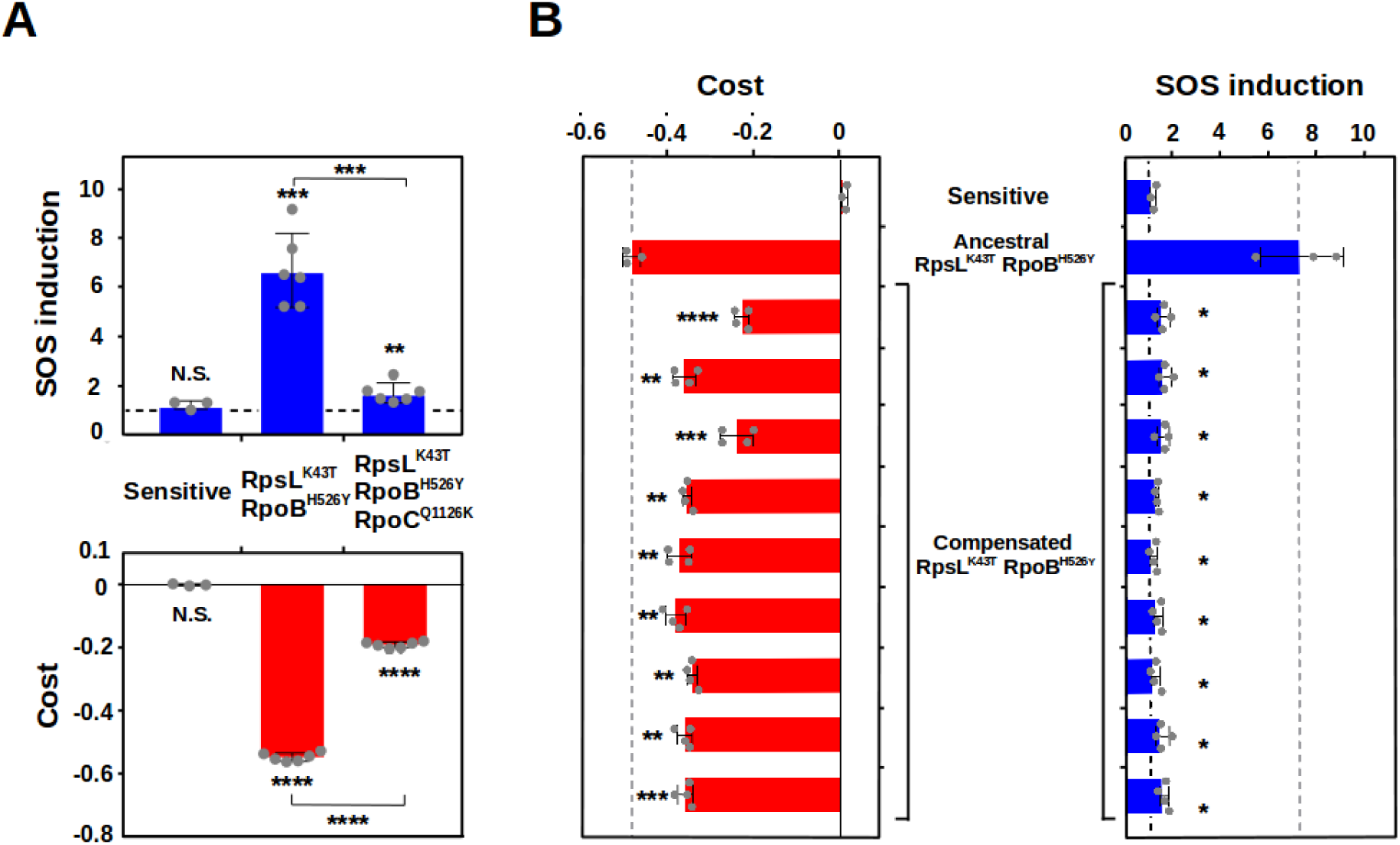
Compensatory evolution cause reduction of DNA breaks. **A.** SOS induction (blue bars) and fitness cost (red bars), of sensitive (left) double resistant (center) and double resistant carrying a compensatory mutation (right) bacteria in LB broth at 4 hours. The dashed line indicates no SOS induction. Error bars represent mean ± standard deviation of independent biological replicates (n≥3). N.S. non-significant; \**P* < 0.05; \*\**P* < 0.01; \*\*\**P* < 0.001; \*\*\*\**P* < 0.0001 (two-tailed Student’s *t* test). **B.** Fitness cost (red bars), and SOS induction (blue bars) of sensitive, ancestral RpsL^K43T^ RpoB^H526Y^ and nine compensated clones in LB broth at 4 hours. The black dashed line indicates no SOS induction. The grey dashed lines mark the cost/SOS of the ancestral double mutant. Error bars represent mean ± standard deviation of independent biological replicates (n≥3). N.S. non-significant; \**P* < 0.05; \*\**P* < 0.01; \*\*\**P* < 0.001; \*\*\*\**P* < 0.0001 (two-tailed Student’s *t* test).

### Increasing transcription-coupled DNA repair reduces the cost of resistance

We then hypothesized that, if DNA breaks contribute to the cost of resistance, enhancing DNA repair should reduce the cost. The RNA polymerase-binding transcription factor DksA has been recently shown to be involved in the transcription-coupled repair of DNA breaks (53, 54). Thus, we hypothesize that overexpression of *dksA* should decrease the cost of resistance. As hypothesized, overproduction of DksA increases the fitness of the double resistant RpsL^K43N^ RpoB^H526Y^, but not that of sensitive bacteria (Figure S2A), indicating that the relationship between DNA breaks and the fitness cost of antibiotic resistance is, indeed, causal.

### Replication speed affects the cost of resistance

Transcription-translation uncoupling can involve increased generation of replication-transcription conflicts, which ultimately generate DNA breaks (42, 55). Replication-transcription conflicts are maximized during fast replication, and less pronounced when bacteria grow slowly (56). Consequently, we hypothesized that the cost of resistance should be expressed mostly when cells are rapidly dividing. Indeed, we observed that, although the costs of Rif^R^ and Str^R^ mutations are similar at 4h and 24h, these costs are generated in the first 4 hours (which comprise exponential growth), while resistance mutations show no cost afterwards, when growth slows down (Figure 3A). We therefore hypothesize that altering replication-transcription conflicts should affect the fitness cost of resistance. To test that hypothesis, we influenced the occurrence of replication-transcription conflitcs through two strategies: negatively by growing our set of mutants in minimal medium, where DNA replication is slower (57), or positively it by overexpressing the DNA replication initiator protein DnaA, which causes simultaneous initiation of multiple replication forks (58). Our hypothesis predicts that DNA breaks should be reduced in minimal medium, where replication-transcription conflicts are less pronounced (Figure 3B). In agreement with our hypothesis, the SOS induction is smaller in minimal than in rich media (Figure 3C, right panel), and the correlation with the cost is much weaker in minimal medium (Figure S2B). Coherently, the number of resistant mutants showing a fitness cost is smaller in minimal medium (8, compared to 13 in rich medium), with four of them even showing a higher fitness than sensitive bacteria (Figure 3C, left panel). Moreover, some of the most costly mutations in rich medium (e. g. RpsL^K43N^ RpoB^H526Y^ and RpsL^K43T^ RpoB^H526Y^) generate a much smaller cost in minimal medium (compare Figure 1A with Figure 3C). Besides this general trend, however, there are resistant mutants (notably, those carrying the allele RpoB^S531F^) that conserve or even increase their fitness cost in minimal medium (compare Figure 1A with Figure 3C), evidencing the existence of additional factors contributing to the cost of resistance (such as the known defects in protein synthesis). Conversely, our hypothesis also anticipates that increasing the occurrence of replication-transcription conflicts by overexpressing DnaA should particularly affect resistant bacteria. As hypothesized, overexpression of *dnaA* severely compromises the viability of double resistant bacteria, whereas sensitive bacteria remains largely unaffected (Figure 3D). Altogether, these results demonstrate the involvement of replication-transcription conflicts in the generation of the cost of antibiotic resistance, also corroborating the key contribution of DNA breaks to it.

**Figure 3.**
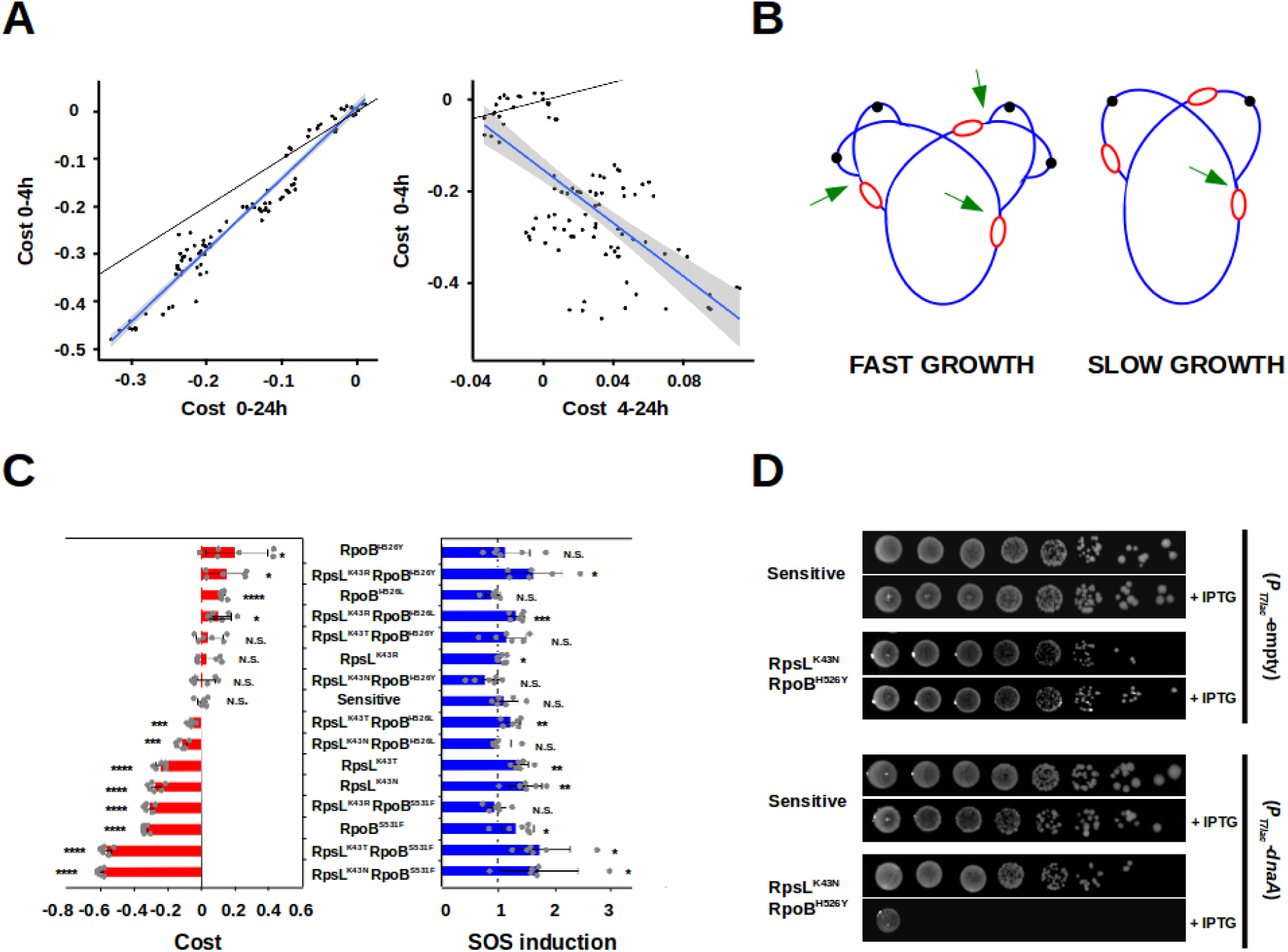
Replication speed affects the cost of resistance. **A.** Left panel: correlation of the fitness cost between time 0 and 4h (y axis) and between time 0 and 24h (y axis), in LB broth. Right panel: correlation of the fitness cost between time 0 and 4h (y axis) and between time 4h and 24h (y axis), in LB broth. Black lines represent the linear regressions if the costs were identical. Blue lines represent the regression lines, and the grey areas represent the 95% confidence intervals. Data from the experiments shown in Figure 1A. **B**. Schematic representation of DNA replication of *E. coli* during fast (left) or slow (right) growth. Black dots represent the origin of replication, and red lines represent transcription forks; green arrows mark regions of potential replication-transcription conflicts. **C.** Fitness cost (red bars), and SOS induction (blue bars) of sensitive bacteria and Str^R^ and/or Rif^R^ mutants in minimal medium at 8 hours. The strains are ordered from lower to higher cost (top to bottom). The dashed line indicates no SOS induction. Error bars represent mean ± standard deviation of independent biological replicates (n=6). N.S. non-significant; \**P* < 0.05; \*\**P* < 0.01; \*\*\**P* < 0.001; \*\*\*\**P* < 0.0001 (two-tailed Student’s *t* test). **D.** Aliquots of serial dilutions (approximately from 5×10^7^ to 5 cells) of sensitive and RpsL^K43N^ RpoB^H526Y^ strains carrying the expression vector pCA24N-gfp, either empty (top, strains RB1317 and RB1321) or harboring the gene *dnaA* (bottom, strains RB1290 and RB1294) and either in the absence or the presence of the inducer IPTG. Each experiment included three biological replicates and three experiments were performed; representative data sets are shown.

### RNAse HI strongly influences the fitness of resistant bacteria

Both transcription-translation uncoupling and replication-transcription conflicts lead to increased formation of R-loops, which cause DNA breaks (42, 59) and exacerbate replication-transcription conflicts themselves by impairing replication fork progression (60). We thus reasoned that depleting RNase HI function, which specifically degrades R-loops (61), should lead to increased DNA breaks and greater costs of Rif^R^ and Str^R^. Accordingly, both DNA breaks and the cost of resistance are greatly exacerbated in the *ΔrnhA* background (Figure 4A, Table 1, Figure S3, Figure S4A). Conversely, mild overproduction of RNase HI can ameliorate both phenotypes in a subset of mutants (Figure S5); however, strong overproduction is toxic for the cell (62, 63), irrespective of its genotype (Figure S6). These results confirm the involvement of R-loops in the uncoupling-mediated production of DNA breaks caused by resistance and suggest that RNase HI can be a target for manipulating the fitness of resistant strains in bacterial populations.

**Figure 4.**
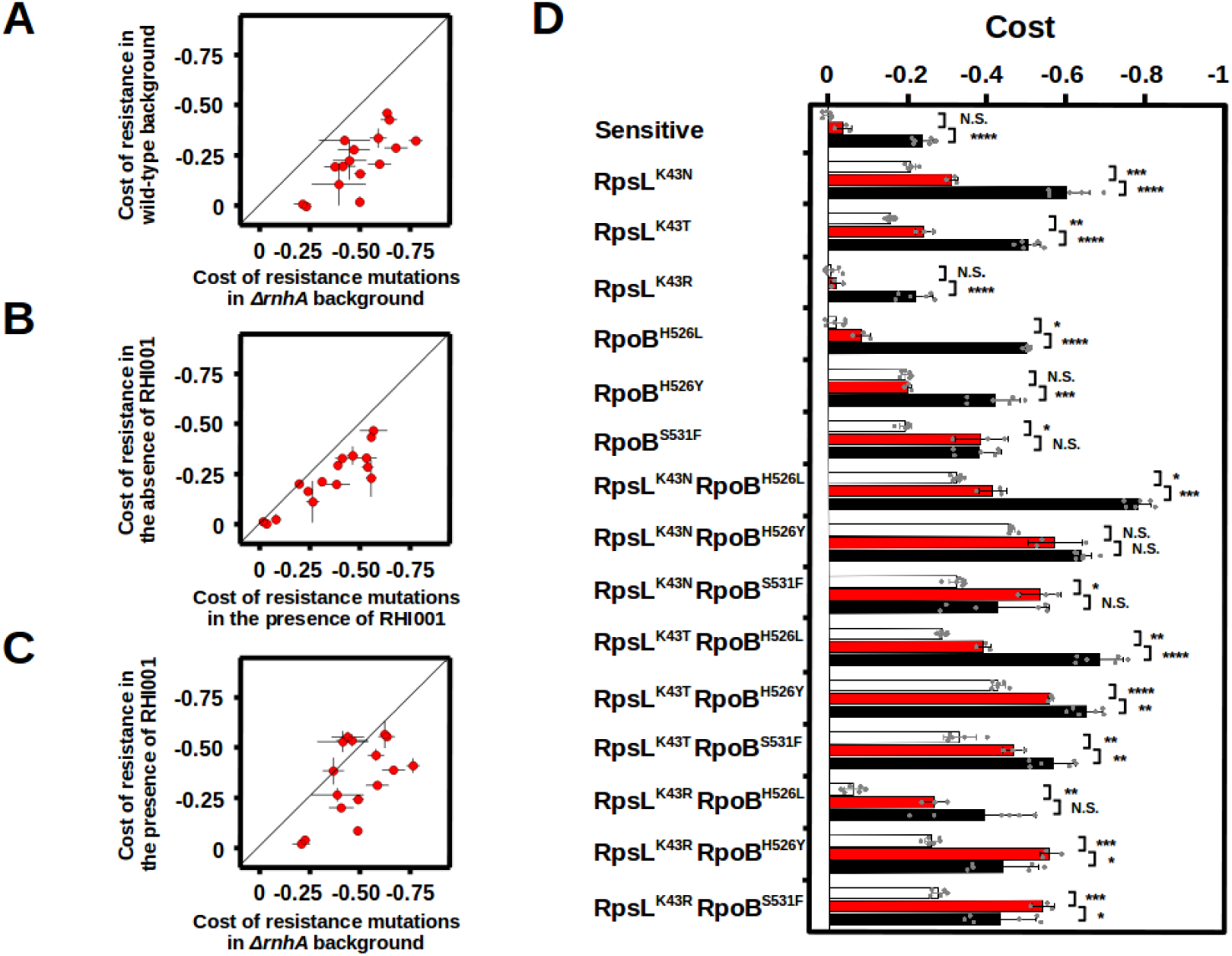
RNAse HI strongly influences the fitness of resistant bacteria. **A.** Correlation between the fitness cost of resistance mutations in wild-type (y axis) or *ΔrnhA* backgrounds (x axis). **B.** Correlation between the fitness cost of resistance mutations in the absence of the RNase HI inhibitor (y axis) or in its presence (x axis). **C.** Correlation between the fitness cost of resistance mutations in the presence of the RHI001 (y axis) or in the *ΔrnhA* background (x axis). The values corresponding to the wild-type background in panel A and to absence of RHI001 in panel B are those shown in Figure 1A. Error bars represent mean ± standard deviation of independent biological replicates (n≥3). The black line in panels A-C represents the linear regression if the costs were identical **D.** Fitness cost of sensitive bacteria and Str^R^ and/or Rif^R^ mutants in the presence of the RNase HI inhibitor RHI001 (red bars). For comparison, the corresponding values in the absence of the inhibitor (data from the experiments shown in Figure 1A) or in a *ΔrnhA* background (data from the experiments shown in panel A and Figure S4A) are represented as white and black bars, respectively. Error bars represent mean ± standard deviation of independent biological replicates (n=3). N.S. non-significant; \**P* < 0.05; \*\**P* < 0.01; \*\*\**P* < 0.001; \*\*\*\**P* < 0.0001 (two-tailed Student’s *t* test). See also Figures S4, S5 and S6.

### RNase HI can serve as a target specific against resistant bacteria

The large effect of lacking RNAse HI function on the fitness of resistant bacteria prompted us to ask whether targeting it could be used to select specifically against resistant bacteria in polymorphic populations. RNase HI inhibitors are currently studied as antiretrovirals (64). A commercially available one, RHI001, has been shown to inhibit the activity of purified *E. coli* RNase HI protein *in vitro* (*65*). In competitions between sensitive and resistant bacteria, we observed that addition of RHI001 to the medium increases the cost of most resistant mutants (Figure 4B), on average by 14%. RHI001 also reduces fitness of double resistant bacteria more than that of sensitive bacteria in the absence of competition (Figure S4B and S4C). Chemical inhibition was not as effective as genetic removal (which increases cost on average by 28%) (Figure 4C and D), as may be expected, since stability, diffusibility across the bacterial envelope, pharmacokinetics and pharmacodynamics of RHI001 *in vivo* are unknown, and potentially suboptimal. Nevertheless, these results suggest that inhibiting RNase HI may be a plausible strategy to select specifically against resistant strains coexisting with sensitive bacteria, as long as resistant strains fail to evolve adaptations that abrogate their extinction. In order to test this hypothesis, we propagated a mixture of sensitive bacteria (CFP-labeled) competing against a pool of five single resistant mutants (YFP-labeled RpsL^K43N^, RpsL^K43T^, RpoB^H526L^, RpoB^H526Y^, and RpoB^S531F^) during 15 days, in the absence of antibiotics. We studied the frequency dynamics of resistant clones under both strong bottlenecks (1:1500 dilutions), where new adaptive mutations are less likely to spread, and weak bottlenecks (1:50 dilutions), where propagation of adapted clones is more probable. In parallel, we performed identical propagations, but in which all strains lack RNase HI, as a proxy for optimal inhibition of RNase HI function. We observed that, in the presence of RNase HI, sensitive bacteria initially outcompete resistant clones but, as the propagation progresses, resistant bacteria increase in frequency - likely due to compensatory evolution - finally reaching coexistence (Figure 5A, blue lines). Remarkably, in the propagations of strains lacking RNase HI, resistant bacteria were completely outcompeted and went extinct by day seven (Figure 5A, red lines) even under mild bottlenecks (Figure 5B). Since the outcome of competitions under laboratory conditions can differ from those within the host (4, 66), we next tested if inhibition of RNase HI would be an effective strategy in an animal model. We used the well-established mouse gut colonization model, which resembles the natural environment of many bacterial pathogens, including *E. coli* (67). We colonized by oral gavage two cohorts of mice (each n=6): one with a 1:9 ratio mixture of CFP-tagged sensitive bacteria and a YFP-labeled double resistant strain [RpsL^K43T^ RpoB^H526Y^, selected based on its ability to efficiently colonize the mammalian gut (68)], and the other with an identical mixture but in which both strains are *ΔrnhA.* We then followed the frequency and absolute numbers [Colony Forming Units (CFUs) per gram of feces] of the resistant strain over time. Remarkably, when RNase HI function is intact, the double resistant strain persists in the gut of all animals at frequencies between 2.3 and 9.3% (Figure 5C, blue lines) and high bacterial loads (Figure 5D, blue lines) two weeks after gavage but, in the absence of RNase HI, resistant clones are rapidly outcompeted by sensitive bacteria (Figure 5C and D, red lines). Indeed, resistant bacteria are driven to extremely low frequencies in as soon as 1 day post-colonization, and their extinction occurs in all the six mice in as soon as 5 days (Figure 5C and D, red lines), demonstrating that RNase HI is a new target for efficiently eliminating resistant strains in the gut. Surprisingly, the absence of RNase HI does not lead to any detectable fitness defect in the sensitive bacteria, as the loads of *E. coli* are similar in the two cohorts of mice (Figure 5E). We then queried if the sequential application of an RNase HI inhibitor followed by an antibiotic could successfully eliminate the entire population of a potential gut pathogenic species composed of sensitive and resistant strains. In this scenario, we hypothesize that an antibiotic treatment should render completely different outcomes depending on the presence of RNase HI function. Indeed, a week-long treatment of streptomycin starting at day 16 after gavage enables resistant clones with intact RNase HI function to rebound and reach high loads (10^7^-10^8^ CFUs/g feces), but it completely erradicates *ΔrnhA* bacteria in all the six mice (Figure 5F). These results demonstrate that RNAse HI function is not only a key determinant of the fitness of resistant bacteria, but also a potential co-adjuvant to eliminate undesirable bacteria in a natural environment for bacterial pathogens such as the mammalian gut. Altogether, these results show the enormous potential of targeting RNase HI as a promising antimicrobial therapy.

**Figure 5.**
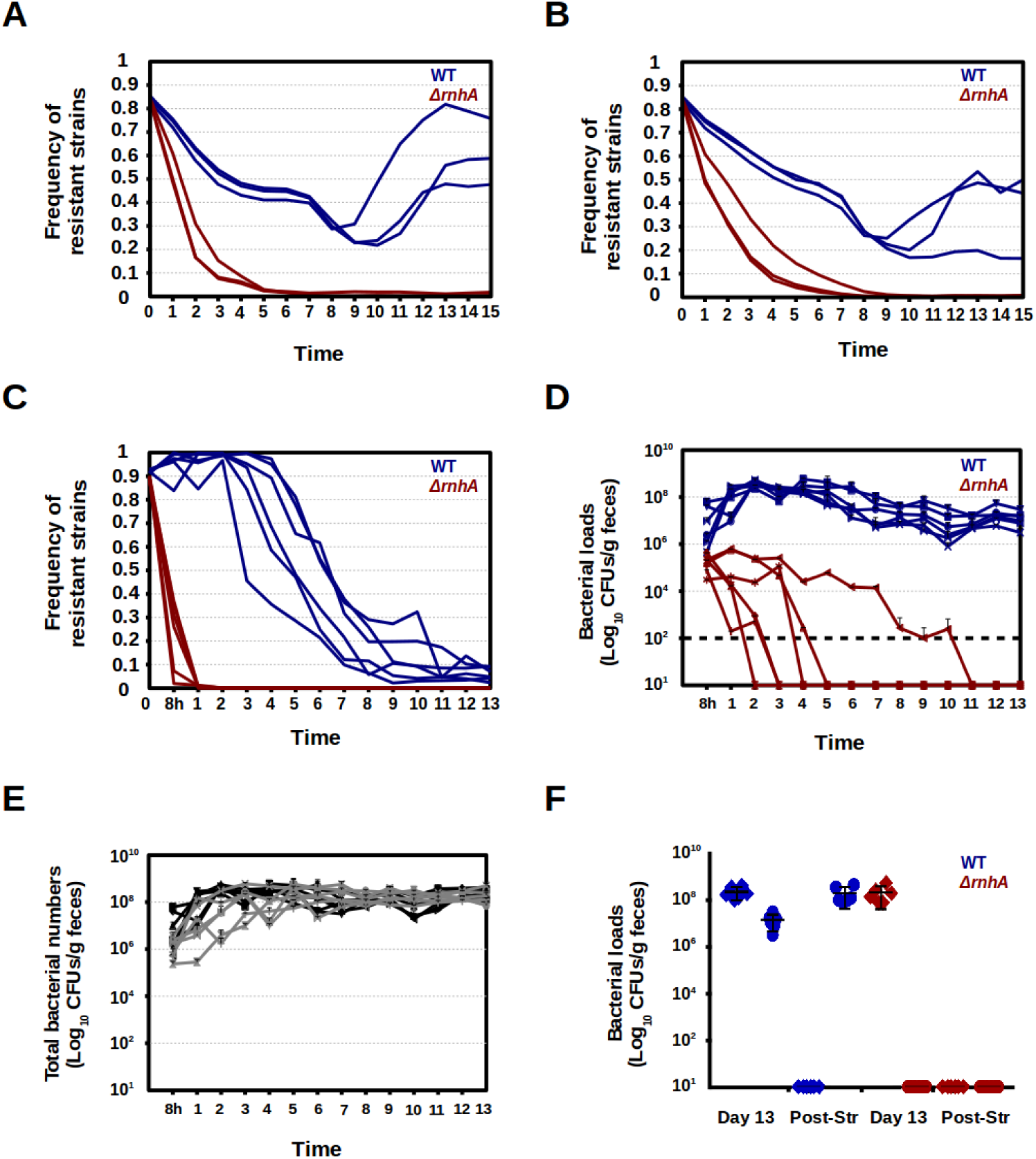
Lack of RNase HI favors outcompetition of resistant mutants by sensitive bacteria. **A and B.** Frequency of single resistant mutants during three independent long-term competitions in rich medium against sensitive bacteria either in a genetic background including RNase HI (blue lines) or in a *ΔrnhA* background (red lines), imposing a strong (1:1500 dilutions, **A**) or a mild (1:50 dilutions, **B**) bottlenecks. **C and D**. Frequency (**C**) and bacterial loads (Log_10_ CFUs/g og feces, **D**) of double resistant mutants during six independent long-term competitions against sensitive bacteria in mice, either in genetic backgrounds including RNase HI (blue lines) or in *ΔrnhA* backgrounds (red lines). Data are represented until extinction was observed in all six animals (13 days in the competitions in *ΔrnhA* backgrounds). The black dashed line represents the limit of detection. **E.** Total (sensitive + double mutant) bacterial loads (Log_10_ CFUs/g of feces) of the strains carrying RNase HI (black lines) or in the *ΔrnhA* background (grey lines) during the competitions in the mouse gut. **F.** Bacterial loads (Log_10_ CFUs/g og feces) at day 13 after gavage and after the week-long streptomycin treatment of sensitive (diamonds) and double resistant bacteria (circles) either in a genetic background carrying RNase HI (blue) or in a *ΔrnhA* background (red). Error bars represent mean ± standard deviation of the results in different mice (n=6).

## Discussion

We show that mutations affecting transcription and translation generate DNA breaks, which explain 73% of the variation in their cost (Figure 1, Table 1, Figure S1). Compensatory evolution indicates that these resistances cause uncoupling between transcription and translation (40), which can result in DNA breaks (42). Indeed, we show that the RpsL^K43N^ RpoB^H526Y^ and RpsL^K43T^ RpoB^H526Y^ double mutants, which combine alleles that cause increased transcription elongation rate (12) and decreased translation rate (26, 69), show the highest costs and DNA breaks (Figure 1). Curiously, inhibition of translation has been shown to alleviate the cost of Rif^R^ mutations in *Pseudomonas aeruginosa* (36). In light of the results presented here, this observation is compatible with a scenario in which Rif^R^ mutations cause a perturbation of transcription-translation coupling by virtue of reduced RNA polymerase processivity, and translation inhibitors promote re-coupling by decreasing ribosomal activity. In agreement with the notion of single resistance mutations causing significant uncoupling, single Rif^R^ or Str^R^ mutations with high cost (70) cause a strong induction of the SOS response (Figure S7). We also showed that RNase HI is a key determinant of the fitness of resistant mutants (Figure 4, Table 1, Figure S3, Figure S4A, Figure S5, Figure S6). Interestingly, transcription-translation uncoupling caused by chemical inhibition of translation can generate R-loops (42,71,72). Thus, perturbations of the coupling between transcription and translation increase the requirement of RNase HI function by bacteria. This opens the possibility to designing novel therapeutic interventions by combining drugs targeting transcription or translation with RNase HI inhibitors in order to enhance the effectiveness of antimicrobial treatments.

We observed that replication-transcription conflicts contribute to the cost of resistance (Figure 3), and that overproduction of DksA, involved in transcription-coupled DNA repair (53, 54), reduces the cost of resistance (Figure S2A). Interestingly, DksA is also involved in preventing replication-transcription conflicts (73). Thus, prevention of the conflicts by DksA might also account for its beneficial effect on resistant bacteria, besides the enhancement of DNA repair. The involvement of replication-transcription conflicts in the cost of resistance is coherent with previous observations, such as the small cost of specific Rif^R^ and/or Str^R^ alleles in minimal medium (24,74,75) or the outcompetition of sensitive bacteria by resistant mutants in aging colonies (76, 77). Interestingly, stable DNA replication (initiation of DNA replication at sites different from *OriC*) can be induced by lack of RNase HI or by conditions that activate SOS (78). Thus, Rif^R^ and Str^R^ mutations, which induce SOS (Figure 1), could additionally favor replication-transcription conflicts via induction of stable DNA replication. This can generate a feed-forward loop of synergistic deleterious effects, which may be further enhanced by the fact that induction of the SOS response causes downregulation of *rnhA* (79). This detrimental runaway loop, as well as the effects of the environment on the cost described above (Figure 1A and Figure 3C), could potentially be exploited therapeutically, via concurrent chemical and dietary treatments designed to synergistically maximize the cost of resistance.

We revealed RNase HI as a promising target specific against resistant bacteria (Figure 4), and showed that lack of RNase HI function favors extinction of resistant clones, preventing compensatory evolution (Figure 5A and B). Importantly, we demonstrated that targeting RNase HI leads to the efficient elimination of resistant bacteria in an animal model as well (Figure 5C and 5D). Interestingly, sensitive bacteria lacking RNase HI do not show fitness defects in the gut (Figure 5E), which hampers the development of resistance to RNase HI inhibition. Moreover, targeting RNase HI prevents recurrence of resistant bacteria upon an antibiotic treatment, favoring the elimination of the entire bacterial population (Figure 5F). These notions are specially important for resistances mediated by mutations (80) or for pathogens that acquire antibiotic resistance exclusively through mutation, such as *Mycobacterium tuberculosis*, which often carries Rif^R^ and Str^R^ mutations (81). Interestingly, RNase HI has been proposed as an antimycobacterial target due to its essentiality in *Mycobacterium smegmatis* (82, 83). The results presented here strongly support the plausibility of this strategy.

An understanding of the determinants of maintenance and dissemination of antibiotic resistance, such as its cost, is urgently needed (1). We show that DNA breaks are key contributors to the cost of resistance and reveal RNase HI as a promising antimicrobial target specific against resistant bacteria, which we validated under laboratory conditions, in the mammalian gut, and by using a repurposed drug. Overall, our results uncover important effects of resistances on bacterial physiology and demonstrate the plausibility of exploiting these effects to develop novel strategies against antibiotic resistance.

## Materials and Methods

### Bacterial strains, media and growth conditions

All the strains used in this study (Data S1) derive from strains RB266, RB323 or RB324, which are derivatives of *E*. *coli* K12 MG1655 (84). Fluorescently labeled strains harbor a copy of either yellow fluorescent protein (YFP) or cyan fluorescent protein (CFP) under the control of the LacI-regulated promoter P_LacO-1_ inserted either in the *yzgL* pseudogene locus or in the *galK* gene, a deletion comprising the entire *lac* operon (to make constitutive the expression of the fluorescent proteins), and the SOS reporter construction *P_sulA_-mCherry* inserted in the *ysaCD* pseudogene locus. The SOS reporter fusion was constructed by replacing a *tetA-sacB* selectable/counterselectable marker (85) located upstream from a *mCherry-FRT-aph-FRT* cassette previously inserted in the *ysaCD* locus by the regulatory regions (150bp upstream from the translation initiation site) of the SOS-regulated gene *sulA*. Resistant mutants additionally carry different chromosomal alleles conferring antibiotic resistance. The fluorescent constructions were generated by Lambda-Red recombineering (86–88), followed by transference to a clean background by P1 transduction (89), and subsequent transduction of the resistance alleles. The clean deletion of *rnhA* was constructed by markerless recombineering using a *tetR-P_tet_-ccdB-cat* selection/counterselection cassette as described by Figueroa-Bossi & Bossi (90), and subsequent transfer of the markerless deletion to a clean background by P1 transduction, using as recipient an isogenic strain carrying a deletion of the nearby gene *proB*, which causes proline auxotrophy, and selecting for growth in minimal medium. The presence of each construction/mutation was assessed by PCR-mediated amplification of the corresponding region and Sanger sequencing. The Gam-GFP construction (49) was generously contributed by Professor Susan M. Rosenberg. The chromosomal construction comprising a copy of the *rnhA* gene under the control of a promoter inducible by arabinose (62) was kindly donated by Dr. Christian J. Rudolph. Derivatives of these strains carrying the *P_sulA_-mCherry* and different resistance mutations were constructed by P1 transduction. The plasmid pRB-5 (carrying the *rnhA* gene under the control of a promoter inducible by anhydrotetracycline) was constructed by PCR amplification of the vector pZS*11 (91) and the construction *tetR-P_LTetO-1_-rnhA* from strain RB1207, subsequent restriction with AatII and HindIII enzymes, ligation and electroporation. The construction *tetR-P_LTetO-1_-rnhA* in strain RB1207 was made by Lambda-Red recombineering (86–88), and selection/counterselection (85), over a previous *P_LtetO-1_-sfGFP* construction (92). The plasmids carrying an inducible copy of either *dksA* or *dnaA* were obtained from the ASKA library of *E. coli* ORF clones (93). Cultures were grown in either Lysogeny Broth (LB, Miller formulation) (94) or M9 broth supplemented with 0.4% glucose (95), in round-bottom 96-well plates incubated at 37°C with shaking (700 r.p.m.) in a Grant-bio PHMP-4 benchtop incubator, unless indicated otherwise. Solid medium was LB containing 1.5% agar. Media were supplemented when necessary with antibiotics at the following concentrations: rifampicin (100 µg/ml), streptomycin (100 µg/ml), ampicillin (100 µg/ml), erythromycin (150 µg/ml), kanamycin (100 µg/ml), chloramphenicol (25 µg/ml).

### Competitive fitness/SOS induction assays

The relative fitness (selection coefficient per generation) of each YFP-tagged resistant strain carrying the SOS reporter was measured by competitive growth against a CFP-labeled isogenic sensitive strain *E*. *coli* K12 MG1655 also carrying the SOS reporter. The formula used to calculate the selection coefficient was *s*=[ln (*NRf* / *NSf*)*−* ln (*NRi* / *NSi*)]ln (*NSf* / *NSi*), being ***NRi*** and ***NRf*** the initial and final number of resistant bacteria, and ***NSi*** and ***NSf*** the initial and final number of sensitive bacteria. The competitor strains were first streaked out of their respective frozen vials, then individual colonies were inoculated separately in medium without antibiotics and incubated overnight (approximately 16h); the next morning, the number of cells in each culture was measured by Flow Cytometry, and 10 µl of 1:1 mixtures of YFP and CFP bacteria were added to 140 µl of medium, at an initial number of approximately 10^6^ cells. The initial and final frequencies of the strains were obtained by counting their cell numbers in the Flow Cytometer. The number of generations was estimated from that of the reference strain (approximately five generations at 4h, and approximately eight generations at 24h), and the selection coefficient was determined as described above for each independent competition. The proportion of SOS-induced bacteria was quantified as the number of either YFP-tagged or CFP-labeled bacteria showing red fluorescence (from the *P_sulA_-mCherry* SOS reporter fusion) above a threshold determined by the fluorescence levels of the control strains *lexA (ind-)* (constitutive repression of the SOS response) and *ΔlexA* (constitutive activation of the SOS response) (96). The data represented is the induction level of each mutant normalized with respect to the induction levels of the sensitive it is competing against. In the competitions including the RNase HI inhibitor 2-[[[3-Bromo-5-(2-furanyl)-7-(trifluoromethyl)pyrazolo[1,5-a]pyrimidin-2-yl]carbonyl]amino]-4,5,6,7-tetrahydro-benzo[b]thiophene-3-carboxylic acid ethyl ester (RHI001, Glixx Laboratories Inc., catalog number GLXC-03982), the medium was supplemented with 500 µM of RHI001, unless indicated otherwise.

### Flow Cytometry

A BD LSR Fortessa^TM^ SORP flow cytometer was used to quantify bacteria, using a 96-well plate High-Throughput Sampler (HTS) and SPHERO fluorescent spheres (AccuCount 2.0 µm blank particles), in order to accurately measure volumes. Bacterial numbers were calculated based on the counts of fluorescently labeled bacteria with respect the known number of beads added to a given volume. The instrument was equipped with a 488 nm laser used for scatter parameters and YFP detection, a 442 nm laser for CFP detection and a 561 nm laser for mCherry detection. Relative to optical configuration, CFP, YPF and mCherry were measured using bandpass filters in the range of 470/20 nm, 540/30 nm and 630/75nm, respectively. The analyzer is also equipped with a forward scatter (FSC) detector in a photomultiplier tube (PMT) to detect bacteria. The samples were acquired using FACSDiVa (version 6.2) software, and analyzed using FlowJo (version 10.0.7r2). All Flow Cytometry experiments were performed at the Flow Cytometry Facility of the Instituto Gulbenkian de Ciência (IGC), Oeiras, Portugal.

### Selection for Erm^R^ bacteria

Fifteen independent colonies of sensitive bacteria were separately inoculated in LB in a 96-well plate, incubated at 37°C with shaking (700 r.p.m) for 7 hours, and 0.1ml of either independent culture was plated onto a LB agar plate supplemented with 150 µg/ml erythromycin, and incubated at 37°C for 5 days (Erm^R^ strains grow in the presence of erythromycin, albeit slowly). Colonies able to grow in these plates were streaked onto plates supplemented with 150 µg/ml erythromycin, in order to further assess their *bona fide* resistance, and the *rplD*, and *rplV* genes of the resistant clones were amplified by PCR and analyzed by Sanger sequencing.

### Microscopy

Early exponential cultures were diluted into pre-warmed medium containing the inducer of the Gam-GFP construction (anhydrotetracycline, 25 ng/ml) and incubated at 37°C with shaking (240 r.p.m.) for 3h, prior to imaging. Bacterial solutions were then placed onto 1% agarose (in 1X PBS) pads mounted in adhesive frames between the microscope slide and a coverglass. Images were acquired on an Applied Precision DeltavisionCORE system, mounted on an Olympus IX71 inverted microscope, coupled to a Cascade II 1024×1024 EM-CCD camera, using an Olympus 100x 1.4NA Uplan SAPO Oil immersion objective, where GFP and mCherry were imaged with FITC (Ex: 475/28, EM: 528/38) and TRITC (Ex: 542/28, Em: 617/73) fluorescence filter sets, respectively, and DIC optics. Images were deconvoluted with Applied Precision’s softWorx software, and prepared for presentation (cropping smaller fields to facilitate visualization, and false-coloring green and red fluorescent signals) using Fiji/ImageJ. All microscopy experiments were performed at the Imaging Facility of the IGC.

### Spot assays

Sensitive and RpsL^K43N^ RpoB^H526Y^ strains carrying the expression vector pCA24N-gfp, either empty or carrying the *dnaA* gene (93) were streaked individually onto LB agar plates supplemented with the appropriate antibiotic (to select for the presence of the plasmid), and incubated overnight at 37°C. The next day, three independent colonies from each strain were inoculated separately in LB broth supplemented with the appropriate antibiotic (150 µl per well) in a 96-well plate and incubated overnight at 37°C with shaking (700 r.p.m). The next day, the OD_600_ of the overnight cultures was measured using a Thermo Scientific Multiskan Go spectrophotometer, bacteria were appropriately diluted in 1X PBS to generate solutions containing approximately 10^10^, 10^9^, 10^8^, 10^7^, 10^6^, 10^5^, 10^4^ or 10^3^ cells/ml. Then 5 µl aliquots of these bacterial solutions (approximately 5×10^7^ to 5 cells) were spotted onto LB agar plates supplemented with the appropriate antibiotic (to select for the presence of the plasmid) and either 0 or 50 µM Isopropyl ß-D-1-thiogalactopyranoside (IPTG) and incubated overnight at 37°C. Each experiment included three biological replicates and three experiments were performed; representative data sets are shown.

### Long-term propagations of polymorphic populations

The CFP-tagged sensitive (either WT or *ΔrnhA*) and the five YFP-labeled resistant bacteria (either RpsL^K43N^, RpsL^K43T^, RpoB^H526L^, RpoB^H526Y^, and RpoB^S531F^ or *ΔrnhA* RpsL^K43N^, *ΔrnhA* RpsL^K43T^, *ΔrnhA* RpoB^H526L^, *ΔrnhA* RpoB^H526Y^, and *ΔrnhA* RpoB^S531F^) were streaked individually onto LB agar plates and incubated overnight at 37°C. The next day, three independent colonies from each strain were inoculated separately in LB broth (150 µl per well) in a 96-well plate and incubated overnight at 37°C with shaking (700 r.p.m). The next day, bacteria were quantified by Flow Cytometry, and 10 µl of 1:1:1:1:1:1 mixtures of the sensitive and either resistant bacteria were added to 140 µl of medium, at an initial number of approximately 10^6^ cells. The initial frequencies of the fluorescent strains were confirmed by Flow Cytometry. Every 24h, during 15 days, bacterial cultures were diluted in 1X PBS by a factor of 10^−2^, then 10 µl of these diluted cultures were transferred to 140 µl fresh LB broth and allowed to grow for additional 24h, reaching approximately 10^9^ cells/ml. In parallel, cell numbers were counted using the Flow Cytometer, in order to measure the frequency of each strain in the mixed population during the experiment, by collecting a sample (10 µl) from the spent culture each day.

### Growth curves

In the experiments involving RHI001, YFP-tagged sensitive and RpsL^K43N^ RpoB^H526Y^ strains were streaked individually onto LB agar plates and incubated overnight at 37°C. The next day, three independent colonies from each strain were inoculated separately in LB broth (150 µl per well) in a 96-well plate and incubated overnight at 37°C with shaking (700 r.p.m). The next day, bacteria were quantified by Flow Cytometry, and approximately 5×10^5^ bacteria were inoculated in 100-well plates containing LB broth supplemented with either 0.5% DMSO (the solvent of RHI001) or 50 µM RHI001 (150 µl per well), and incubated at 37°C with continuous shaking (medium amplitude, duration 5s, 10s interval, and stopping 5s before each measurement) in a Bioscreen C (Oy Growth Curves Ab Ltd.) benchtop microplate reader, measuring OD_600_ every 30 minutes during 12h. In the experiments involving sensitive and RpsL^K43N^ RpoB^H526Y^ strains carrying the expression vector pCA24N-gfp, either empty or carrying the gene *dksA* (93), the strains were streaked individually onto LB agar plates supplemented with the appropriate antibiotic (to select for the presence of the plasmid), and incubated overnight at 37°C. The next day, three independent colonies from each strain were inoculated separately in LB broth supplemented with the appropriate antibiotic (150 µl per well) in a 96-well plate and incubated overnight at 37°C with shaking (700 r.p.m). The next day, the OD_600_ of the overnight cultures was measured using a Thermo Scientific Multiskan Go spectrophotometer, bacteria were appropriately diluted in 1X PBS, and approximately 5×10^5^ bacteria were inoculated in flat-bottom 96-well plates containing LB broth (150 µl per well) supplemented with the appropriate antibiotic and either 0 or 50 µM IPTG, and incubated at 37°C (1°C up to bottom gradient, “lid on” option) with double orbital shaking (set as “fast”) in a Biotek SYNERGY H1 benchtop microplate reader, measuring OD_600_ every 30 minutes during 24h.

### Animal experiments

Adult C57BL/6J female mice (six to eight weeks old) were kept in ventilated cages under specified pathogen free (SPF) barrier conditions at the animal facility of the IGC. To overcome bacterial colonization resistance, mice were treated with streptomycin (5 g/L) in drinking water during seven days and then put on regular autoclaved water for two days to wash out the antibiotic from the intestine. The following day, animals were deprived of food and water during the 4 hours prior to inoculation by oral gavage of 100 µl of bacterial suspension containing ∼10^9^ cells. All animals were then individually-caged, with food and water being available *ad libitum*. Each animal (6 per group) was gavaged with a mix (9:1) of CFP-tagged sensitive (either WT or *ΔrnhA*) and YFP-labeled double resistant bacteria (either RpsL^K43T^ RpoB^H526Y^ or *ΔrnhA* RpsL^K43T^ RpoB^H526Y^). In order to obtain the bacterial suspensions, the strains were streaked individually onto appropriate antibiotic-supplemented LB agar plates and incubated overnight at 37°C. The next day, three independent colonies from each strain were inoculated separately in assay tubes containing 5ml of BHI broth supplemented with the appropriate antibiotic and bacterial cultures were grown overnight at 37°C with shaking (240 r.p.m). The next day, each bacterial culture was diluted 100-fold (20 µl of overnight culture in 2 ml of fresh BHI broth, in 125 ml flasks) and incubated at 37°C with shaking (240 r.p.m) until reaching an OD_600_ ≈ 2. Then the cultures were washed once with 1X PBS and Flow cytometry was used to assess the appropriate cells numbers and prepare bacterial mixes for gavage. Mouse fecal pellets were collected at 8 hours after gavage, and afterwards on a daily basis during 13 days, to analyze the loads and frequency of each strain colonizing the gut along time. Fecal samples were stored in 15% glycerol at −80°C. In order to measure the loads (CFUs/g of feces) and the frequency of each strain along the experiment, fecal samples were plated on appropriate antibiotic-supplemented plates and incubated overnight to assess CFUs of CFP- or YFP-labeled bacteria using a SteREOLumar Carl Zeiss fluorescent stereoscope. Every strain that remained below the detection limit (∼10^2^ CFUs/g of feces) for three consecutive time points was considered extinct. For the post-gavage antibiotic treatment, streptomycin (5 g/L) was dispensed in drinking water during seven days. The animal experiments included in this work were reviewed and approved by both the Ethics Committee and the Animal Welfare Body of the IGC (license A009.2018) and by the Direção Geral de Alimentação e Veterinária (DGAV, Portuguese national entity that regulates the use of laboratory animals; license 009676). All experiments conducted on animals followed the Portuguese (Decreto-Lei n° 113/2013) and European (Directive 2010/63/EU) legislations concerning housing, husbandry and animal welfare.

### Statistical analyses

All analyses were conducted using Libreoffice Calc version 5.4.1.2 (www.libreoffice.org) software, R version 3.4.4 (www.r-project.org) software via RStudio version 1.1.442 interface (www.rstudio.com) and GraphPad Prism version 7.04 (www.graphpad.com). For each set of competitions (n≥3), two-tailed unpaired Student’s *t*-tests were used. For testing the association between the fitness cost and the level of SOS induction, a linear regression of the selection coefficient (s) with the logarithm of the fold change in SOS induction of each resistant clone (with respect to the sensitive bacteria it is competing against in the same well), was used.

## Acknowledgements

The authors thank Professor Susan M. Rosenberg and Dr. Christian J. Rudolph for kindly providing bacterial strains, the Flow Cytometry and Advanced Imaging facilities of Instituto Gulbenkian de Ciência for technical assistance, Miguel Godinho for useful discussions, and Jonathan Howard, Karina Xavier, Leonardo Gastón Guilgur, Pol Nadal Jiménez and the members of the Gordo and Xavier labs for critically reading earlier versions of this manuscript. This work was funded by the the Marie Sklodowska-Curie Actions (MSCA) with the fellowship 746690-ResistEpist-H2020-MSCA-IF-2016/H2020-MSCA-IF-2016, to R.B., and partially supported by the PREPARE project (JPIAMR/0001/2016-ERA NET), and by ONEIDA and Congento projects (LISBOA-01-0145-FEDER-016417 and LISBOA-01-0145-FEDER-022170), both co-funded by FEEI - “Fundos Europeus Estruturais e de Investimento” from “Programa Operacional Regional Lisboa 2020”, and by national funds from “Fundação para a Ciência e a Tecnologia” (FCT), to I.G. R.B. was also supported by the FCT with the fellowship SFRH/BDP/109517/2015. The authors declare no conflicts of interest.

## Author contributions

R.B: conceptualization, formal analysis, funding acquisition, investigation, methodology, visualization, writing – original draft. N.F: formal analysis, investigation, methodology, writing – review & editing. I.G: conceptualization, formal analysis, funding acquisition, methodology, project administration, resources, supervision, writing – review & editing.

## Data availability

All the strains used (Data S1) are available via Material Transfer Agreement (MTA). The raw data of the experiments shown in all figures and the table are available in Data S2. The images analyzed to obtain the data shown in Table 1, and those used in Figures S1 and S3 are available in the public data repository Zenodo (DOI: 10.5281/zenodo.3381746).

## Supplemental Figures

**Figure S1.**
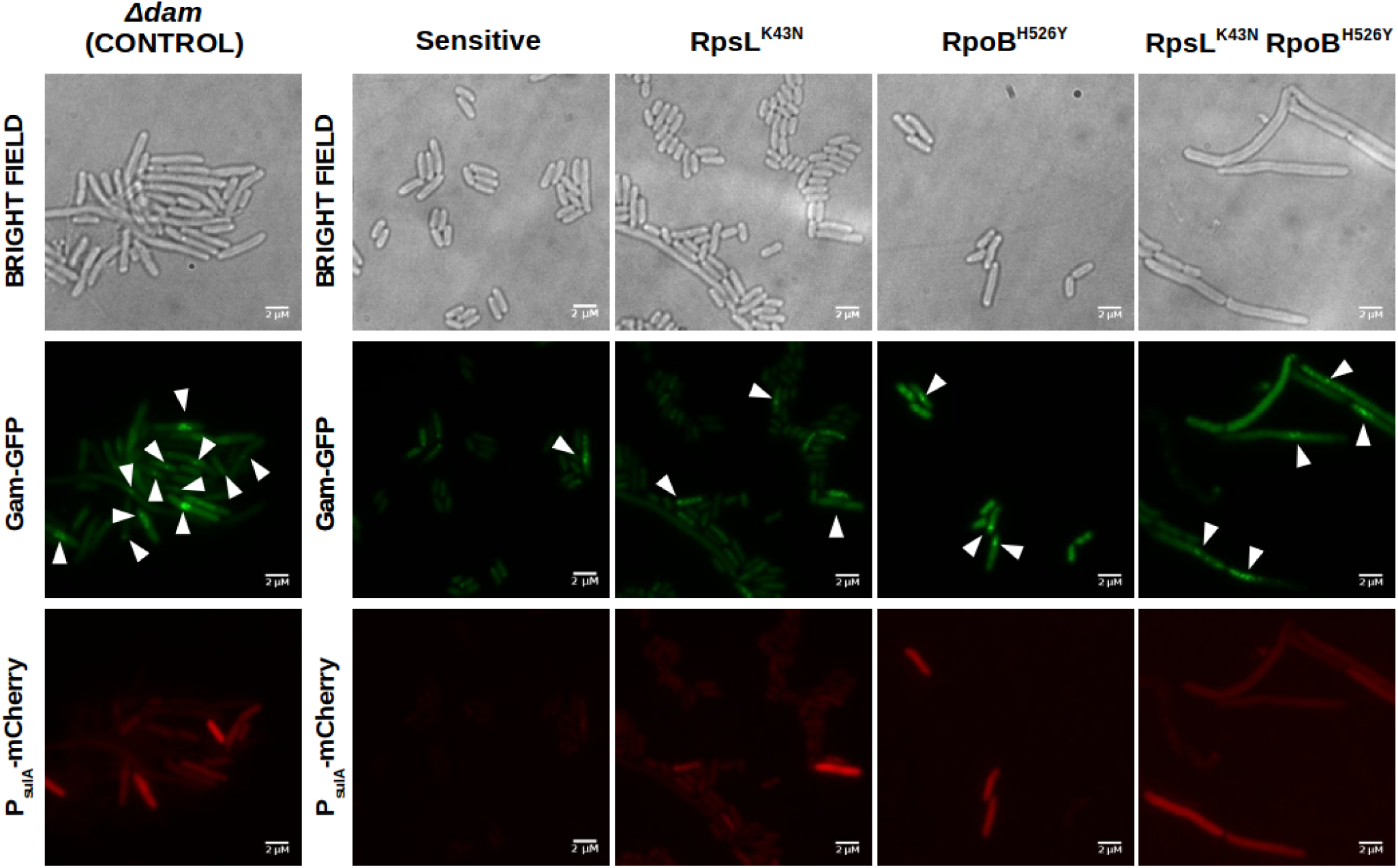
Visualization of DNA breaks and SOS induction of sensitive, single and double resistant strains by epifluorescence microscopy. Bacterial cells show a disperse faint green fluorescence distributed along the cytoplasm, unless DNA breaks are present; upon generation of DNA breaks, the disperse fluorescence concentrates in bright fluorescent foci (central panels, false-colored in green). Red fluorescence (bottom panels, false-colored in red) is absent until the SOS response is induced due to the presence of DNA breaks. Cell elongation (top panels) is a well-known phenotype derived from the inhibition of cell division by SOS-regulated proteins.

**Figure S2.**
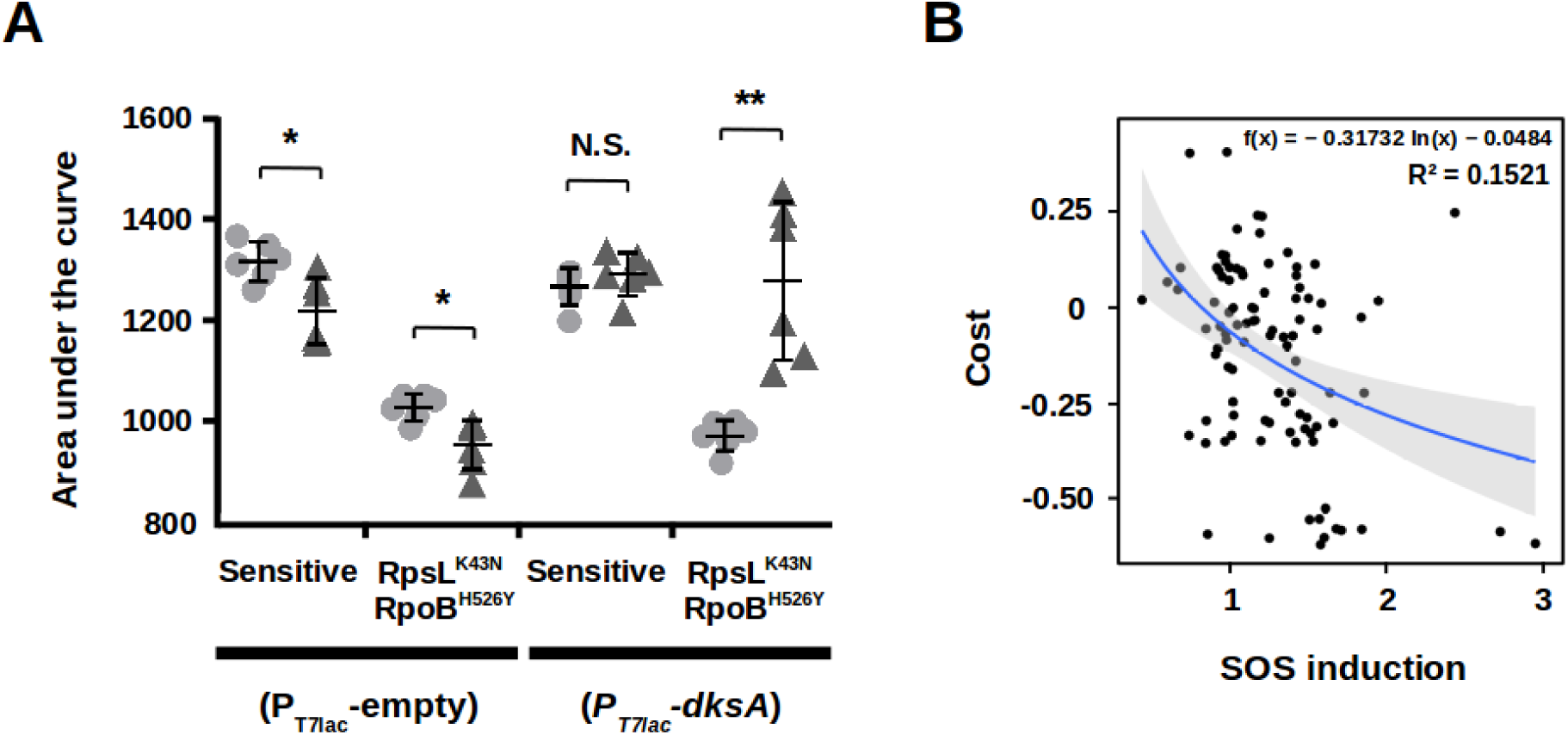
**A**. Area under the curve (24h in LB broth) of sensitive and double resistant bacteria harboring an expression vector either empty (left) or carrying the *dksA* gene in the absence (circles) or presence (triangles) of the inducer. Error bars represent mean ± standard deviation of independent biological replicates (n =6). N.S. non-significant; \**P* < 0.05; \*\**P* < 0.01; \*\*\**P* < 0.001; \*\*\*\**P* < 0.0001 (two-tailed Student’s *t* test). **B**. Correlation between the fitness cost (y axis) and the SOS induction (x axis) at 8h in M9 broth supplemented with 0.4% glucose (data represented in figure 3C). The blue line represents the logarithmic regression line, and the grey area represents the 95% confidence interval.

**Figure S3.**
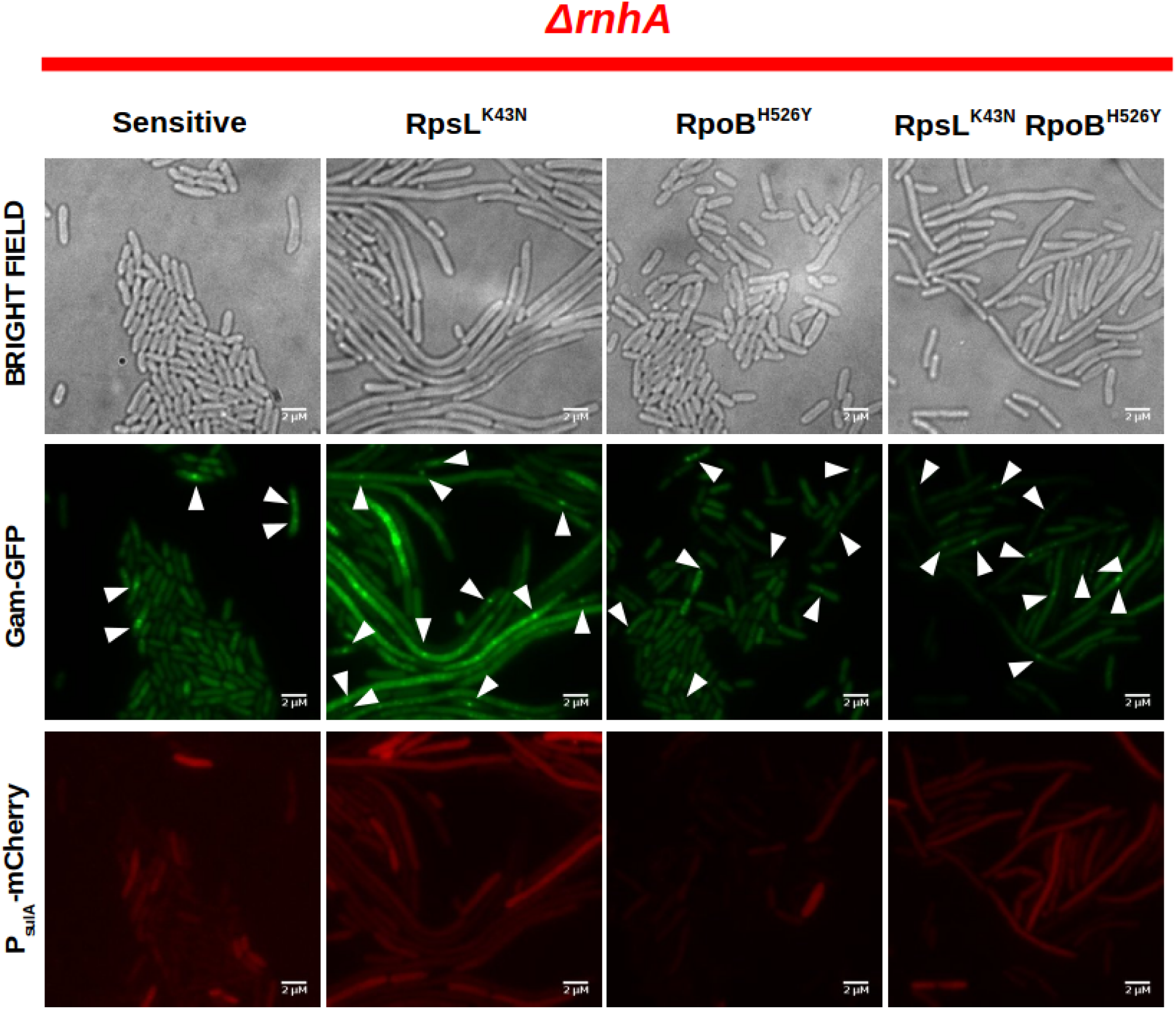
Visualization of DNA breaks and SOS induction of sensitive, single and double resistant strains in *ΔrnhA* background by epifluorescence microscopy. Bacterial cells show a disperse faint green fluorescence distributed along the cytoplasm, unless DNA breaks are present; upon generation of DNA breaks, the disperse fluorescence concentrates in bright fluorescent foci (central panels, false-colored in green). Red fluorescence (bottom panels, false-colored in red) is absent until the SOS response is induced due to the presence of DNA breaks. Cell elongation (top panels) is a well known phenotype derived from the inhibition of cell division by SOS-regulated proteins.

**Figure S4.**
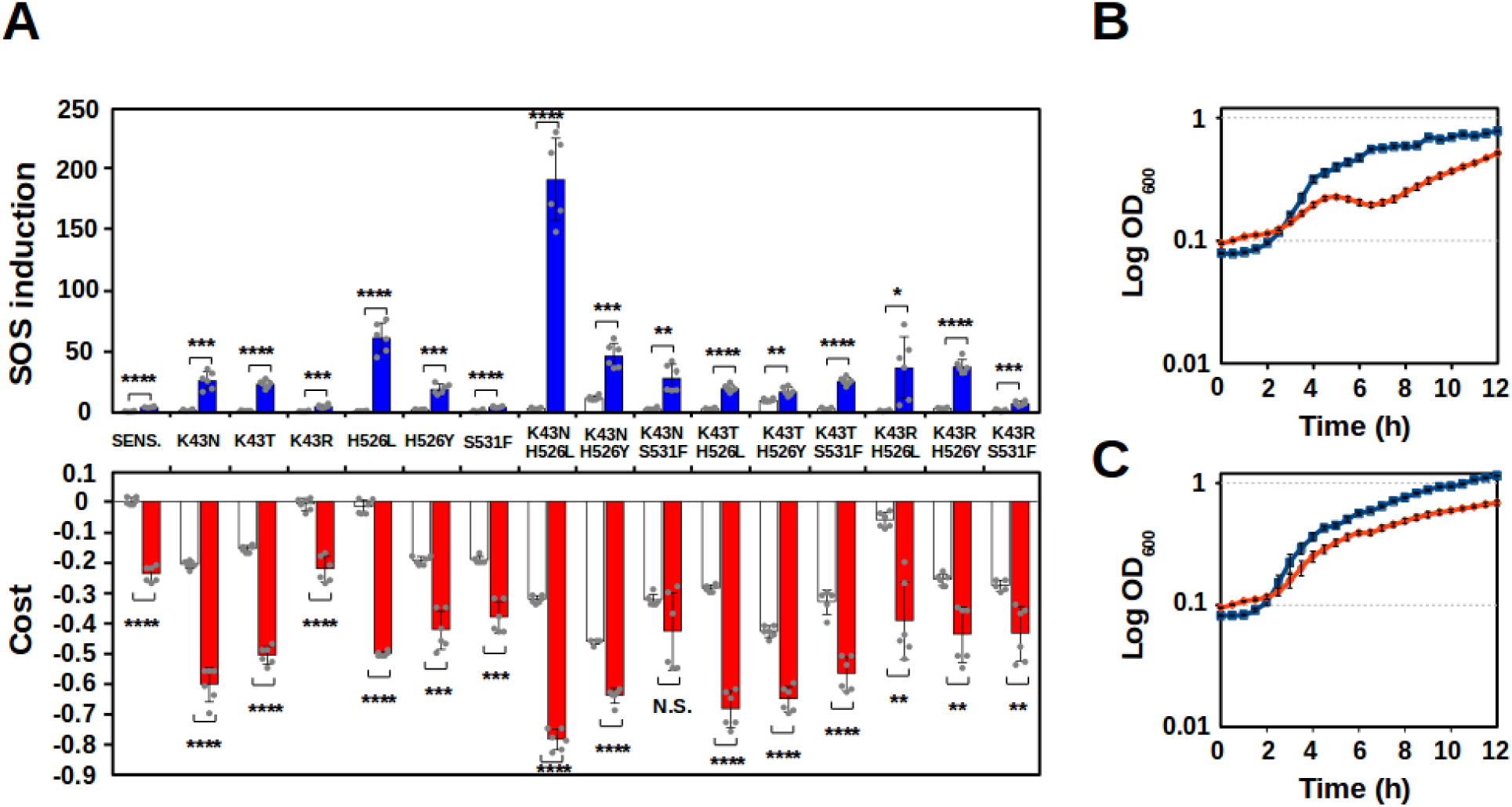
Depletion of RNase HI increases fitness cost and SOS induction. **A.** SOS induction (blue bars) and fitness cost (red bars), of sensitive bacteria and Str^R^ and/or Rif^R^ mutants (labels represent the corresponding allele/s) in a *ΔrnhA* background. White bars represent the corresponding values in bacteria with an intact *rnhA* gene (data from the experiments shown in Figure 1A), for comparison. Error bars represent mean ± standard deviation of independent biological replicates (n=6). N.S. non-significant; \**P* < 0.05; \*\**P* < 0.01; \*\*\**P* < 0.001; \*\*\*\**P* < 0.0001 (two-tailed Student’s *t* test). **B.** Growth curves of RpsL^K43N^ RpoB^H526Y^ double mutant in the presence of either DMSO (blue lines) or RHI001 (orange lines). Points and error bars represent mean ± standard deviation of independent biological replicates (n=3). **C.** Growth curves of sensitive bacteria in the presence of either DMSO (blue lines) or RHI001 (orange lines). Points and error bars represent mean ± standard deviation of independent biological replicates (n=3).

**Figure S5.**
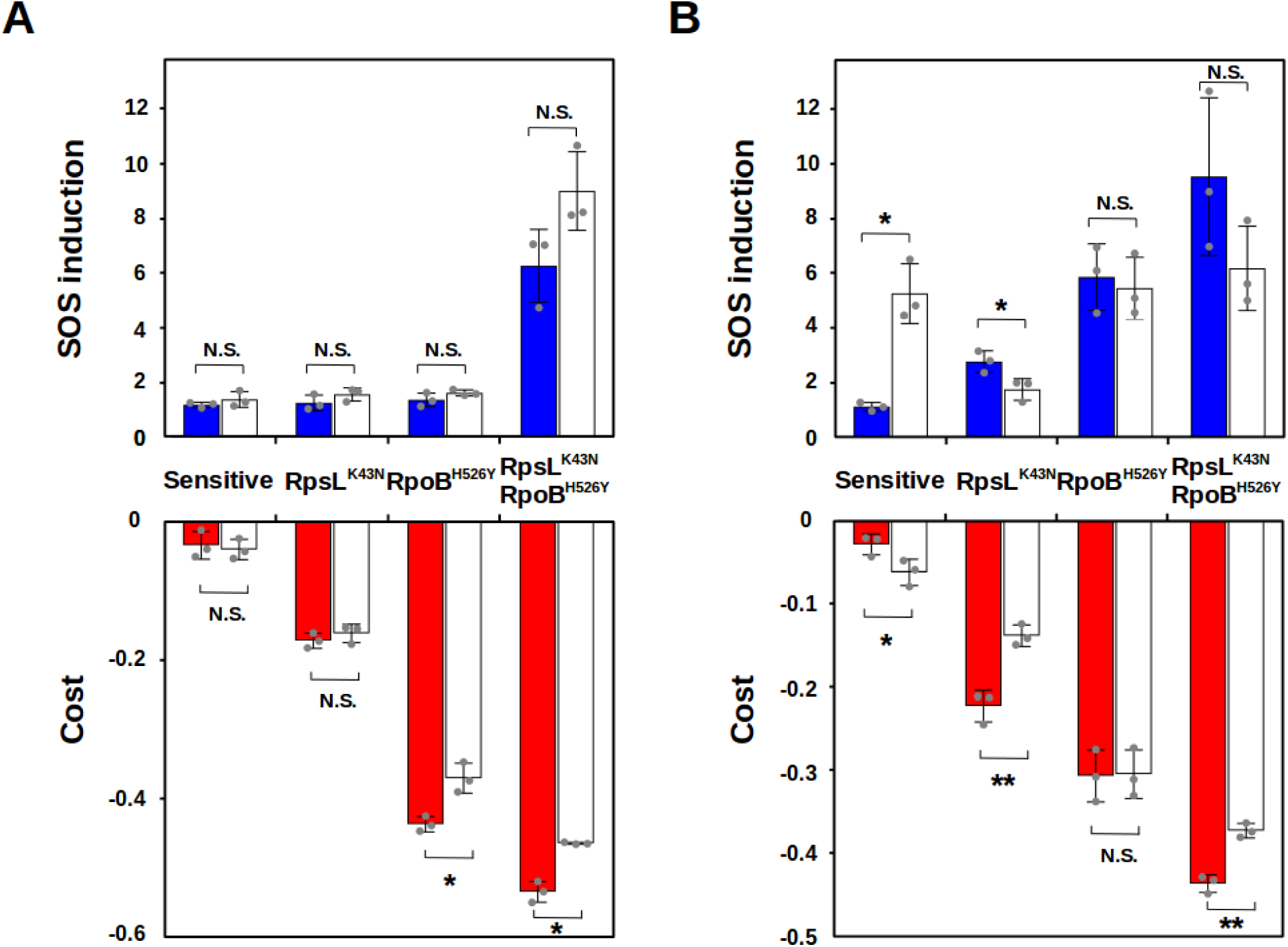
Mild overproduction of RNase HI can ameliorate both cost and SOS induction. SOS induction (top) and fitness cost (bottom), of sensitive bacteria (left), two single resistant mutants (center) and the double mutant combining these two alleles (right) in LB broth at 4h (**A**) or 24h (**B**) in a background harboring an additional chromosomal copy of the gene encoding the RNase HI (*rnhA*) under the control of the arabinose promoter, either in the absence (red bars) or the presence (white bars) of arabinose. Error bars represent mean ± standard deviation of independent biological replicates (n=3). N.S. non-significant; \**P* < 0.05; \*\**P* < 0.01; \*\*\**P* < 0.001; \*\*\*\**P* < 0.0001 (two-tailed Student’s *t* test).

**Figure S6.**
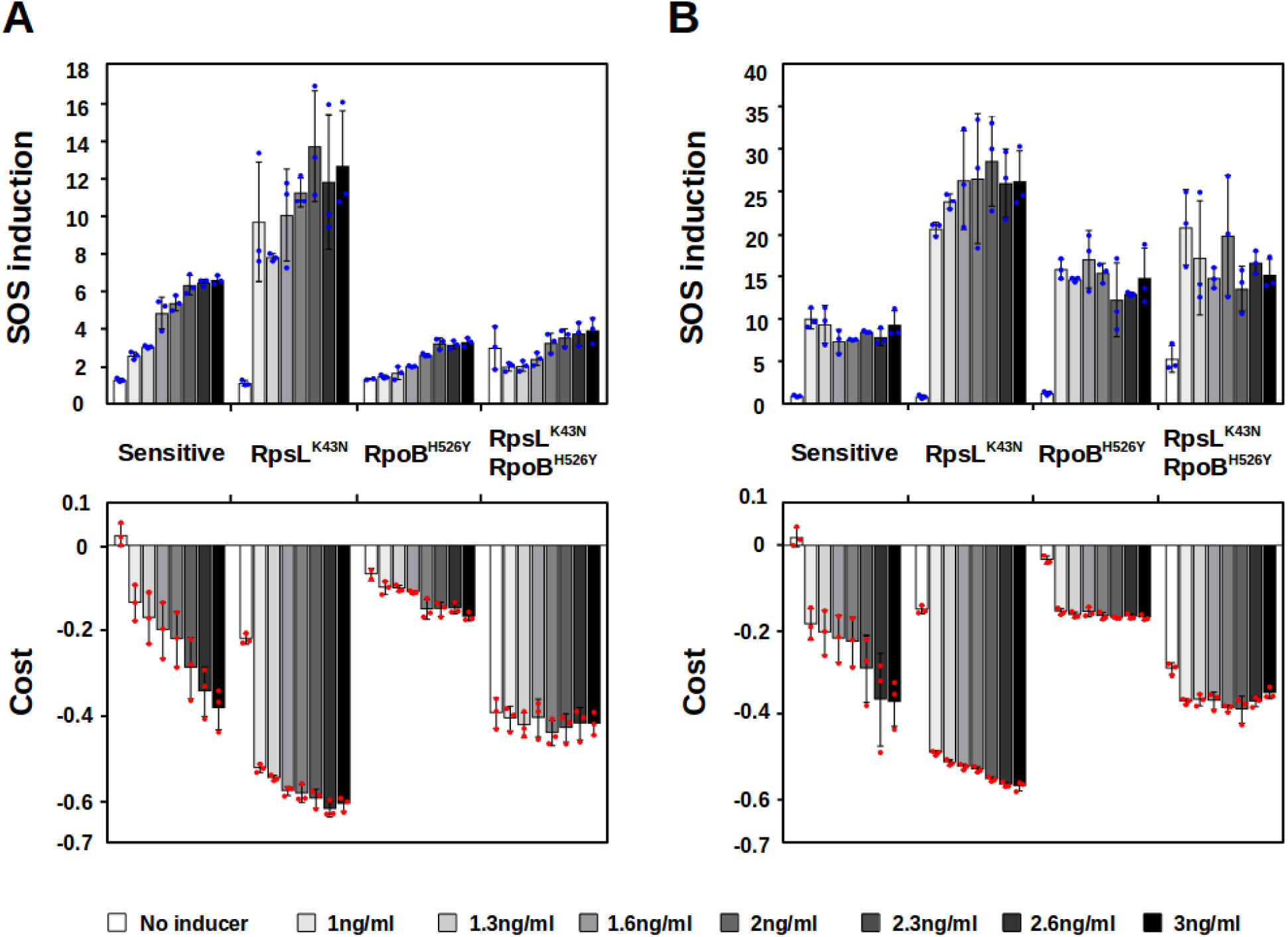
Strong overproduction of RNase HI increases cost and SOS induction in all the backgrounds. SOS induction (top) and fitness cost (bottom), of sensitive bacteria (left), two single resistant mutants (center) and the double mutant combining these two alleles (right) in LB broth at 4h (**A**) or 24h (**B**) in a background harboring a plasmid carrying a copy of the RNase HI gene (*rnhA*) under the control of a promoter inducible by anhydrotetracycline, either in the absence (white bars) or the presence of different concentrations of the inducer (greyscale, from light grey to black bars). Error bars represent mean ± standard deviation of independent biological replicates (n≥2). N.S. non-significant; \**P* < 0.05; \*\**P* < 0.01; \*\*\**P* < 0.001; \*\*\*\**P* < 0.0001 (two-tailed Student’s *t* test).

**Figure S7.**
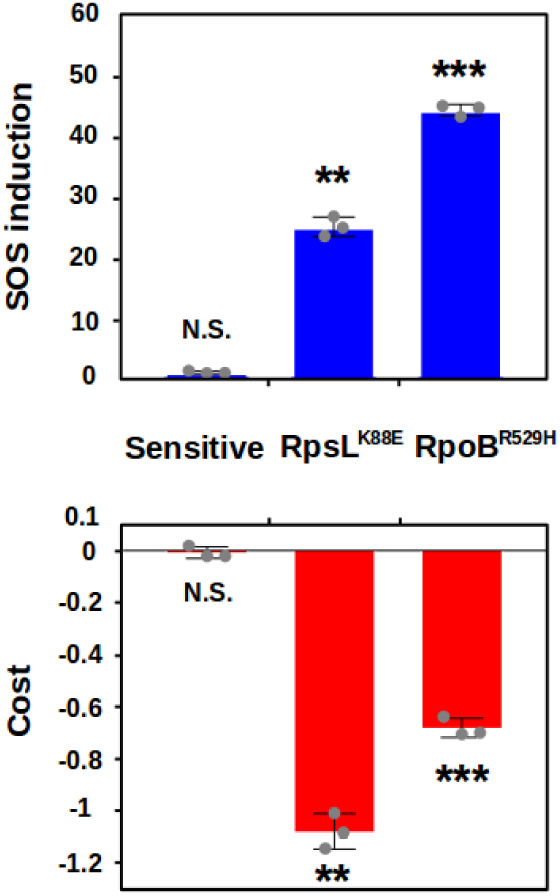
Costly single mutants show increased SOS induction. SOS induction (blue bars) and fitness cost (red bars), of sensitive bacteria (left) and two costly single resistant mutants (center and right). Error bars represent mean ± standard deviation of independent biological replicates (n=3). N.S. non-significant; \**P* < 0.05; \*\**P* < 0.01; \*\*\**P* < 0.001; \*\*\*\**P* < 0.0001 (two-tailed Student’s *t* test).

